# AmpUMI: Design and analysis of unique molecular identifiers for deep amplicon sequencing

**DOI:** 10.1101/288118

**Authors:** Kendell Clement, Rick Farouni, Daniel E. Bauer, Luca Pinello

## Abstract

**Motivation:** Unique molecular identifiers (UMIs) are added to DNA fragments before PCR amplification to discriminate between alleles arising from the same genomic locus and sequencing reads produced by PCR amplification. While computational methods have been developed to take into account UMI information in genome-wide and single-cell sequencing studies, they are not designed for modern amplicon based sequencing experiments, especially in cases of high allelic diversity. Importantly, no guidelines are provided for the design of optimal UMI length for amplicon-based sequencing experiments.

**Results:** Based on the total number of DNA fragments and the distribution of allele frequencies, we present a model for the determination of the minimum UMI length required to prevent UMI collisions and reduce allelic distortion. We also introduce a user-friendly software tool called AmpUMI to assist in the design and the analysis of UMI-based amplicon sequencing studies. AmpUMI provides quality control metrics on frequency and quality of UMIs, and trims and deduplicates amplicon sequences with user specified parameters for use in downstream analysis. AmpUMI is open-source and freely available at http://github.com/pinellolab/AmpUMI.

## 1 Introduction

Next-generation sequencing technologies have enabled the rapid and cost-effective translation of biological DNA or RNA sequences to short sequencing reads that can be used to analyze and understand the genome. In most library preparation protocols, genomic material must be amplified using PCR to ensure successful sequencing. Exponential amplification is sequence dependent and differences in amplification rate may arise from variation in sequence composition (Aird*et al*., 2011). Duplicate sequencing reads resulting from PCR amplification may lead to biases in results or incorrect conclusions about the actual frequency of that read.

Many approaches have been developed for the deduplication of next-generation reads from across the genome. Some tools such as FastUniq (Xu *et al*., 2012) or Fulcrum (Burriesci *et al*., 2012) filter raw sequencing output in FASTQ format and remove any reads with the same sequence. Commonly used tools such as Picard MarkDuplicates (Li*et al*., 2009), samtools rmdup (Li, 2011), and SEAL (Pireddu *et al*., 2011) align the reads to the genome, and identify reads aligning to the same genomic position as duplicates. However, if two DNA fragments from different cells produce reads aligning to the same location, they will be incorrectly called as PCR duplicates, even though they originated from two different molecules. As sequencing depth increases, so does the chance of finding apparently duplicate reads that align to the same genomic location but are from different DNA fragments. In addition, nonuniform genomic fragment formation (e.g., short exons, restriction enzyme digestion, etc) can also increase the apparent rate of PCR duplicates if sequence identity and alignment coordinates are the only criteria for identifying PCR duplicates.

Although PCR deduplication may not affect experimental outcome in cases of calling reference variants from genome wide sequencing (Ebbert *et al*., 2016), library preparation strategies that rely on many PCR amplification cycles (such as single-cell experiments) are susceptible to errors introduced by amplification bias and should be corrected (Islam *et al*., 2014). In libraries with potentially low sequence complexity, PCR amplification bias may go undetected and lead to inaccurate interpretation of sequencing results. For example, accurate allele quantification may be distorted when identifying rare cancer mutations (Kinde *et al*., 2011; Kukita *et al*., 2015; Mansukhani *et al*., 2017) or when measuring the genomic outcomes of CRISPR genome editing experiments (Pinello *et al*., 2016) through amplicon sequencing of a specific target region.

In order to address the problem of duplicate reads from a biological perspective, library generation protocols have been developed that tag DNA fragments with a sequence of randomly selected nucleotides or partially degenerate nucleotides, called a unique molecular index (UMI). After sequencing, reads arising from PCR duplicates will all have the same UMI and all but one of these reads are marked as duplicates as shown in Figure 1a. In downstream analysis these reads are normally ignored, but they can also be used in error correction or in estimating sequencing error rates.

**Fig. 1.**
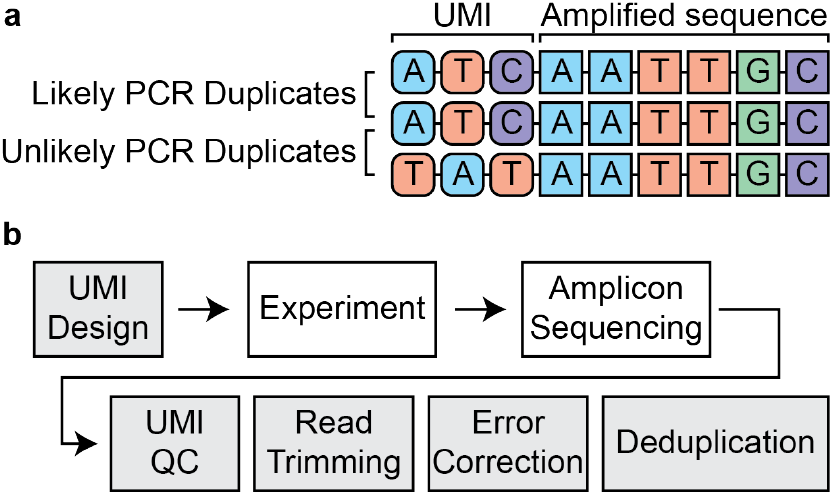
a. Schematic showing utility of UMIs in identifying PCR duplicates. In libraries using UMIs, a short sequence of random nucleotides is added to each DNA fragment before PCR amplification. All PCR products of that read will contain the same UMI. After library sequencing, DNA fragments with the same sequence (shown as square nucleotides on the right part of the read) can be identified as either PCR duplicates or not, based on the UMI sequence (shown as rounded nucleotides on the left part of the read). b. Outline of a standard experiment utilizing UMI technology. The steps shown in gray are computational processing steps performed starting with raw FASTQ output from amplicon sequencing, and are the procedures performed by our software, AmpUMI.

Although there are many benefits to using UMIs, there is a dearth of documented design rationale for selecting sufficient UMI length, and no existing tools to aid experimental designers who may not feel comfortable addressing the probabilities of UMI collision. In one case, a UMI length is chosen so that the number of unique UMIs ‘far exceeds’ the number of primer molecules used in the experiment (Kou *et al*., 2016). However, this may lead to overestimation of the required UMI length.

Several tools to analyze UMI datasets are available, but most are designed for genome-wide sequencing analysis, and require aligned sequence files in bam format as input. UMI-Reducer (Mangul *et al*., 2017), Je (Girardot *et al*., 2016), Debarcer (Stahlberg *et al*., 2016), and UMI-tools (Smith *et al*., 2017) are tools that take UMI information into account in the deduplication process, but are designed for genome-wide sequencing analyses. These methods first trim the UMI and associated sequencing adapters from the sequenced read, and append the UMI to the read name. Next, the trimmed sequencing reads are mapped to the genome. Finally, each read is scanned to identify other duplicate reads that align to the same genomic location and have the same UMI. However, these existing algorithms and methods are not well-suited to amplicon sequencing, where every read aligns to same genomic position. pRESTO (Vander Heiden *et al*., 2014) and MAGERI (Shugay *et al*., 2017) have the capability to work on raw FASTQ files, and can collapse reads by unique UMIs. However, the UMI parsing capabilities for both tools are rigid and, for example, do not allow for degenerate bases in the UMI definition.

Additionally, existing packages attempt to minimize nucleotide variation due to sequencing error by merging reads with one or two differences between the reads (Xu *et al*., 2017). However, in some amplicon experiments, particularly those attempting to identify rare genomic variants at a locus, merging reads with small differences could obscure these rare variants. To address the problem of identifying sequencing errors, an alternate sequencing approach amplifies DNA fragments in many rounds of PCR, and then only considers sequencing reads appearing multiple times for downstream analysis (Stahlberg *et al*., 2017).

Our approach overcomes these challenges by providing users an end-to-end software solution for amplicon sequencing experiments called AmpUMI. AmpUMI aids in both UMI design and subsequent processing of sequencing reads to perform error correction and deduplication. (Figure 1b).

## 2 Approach

### 2.1 Estimating minimum UMI length requirements

Careful UMI design should be performed in the planning stages of an experiment, especially since it is not easy to measure the deleterious effects of inadequate UMI design after sequencing by examining the reads.

One of the most common problems is UMI collision, defined as the event of observing two reads with the same sequence and same UMI barcode but originating from two different genomic molecules. UMI collision is a function of the number of UMIs used, the number of unique alleles, and the frequency of each allele in the population. In genome-wide sequencing experiments, the chance of UMI collision is very low because the number of reads sharing the same sequence is very small. In this study, we focus on amplicon sequencing, in which a specific location in the genome is sequenced in many different cells, usually for the purpose of identifying or quantifying rare alleles. In this case, the sequencing depth is much greater than genome-wide sequencing and many alleles from different genomic molecules will share the same sequence. Because of this, the possibility of UMI collisions is much higher and needs to be taken into consideration in UMI design and analysis of sequenced reads.

In amplicon sequencing studies, it may be tempting to design conservatively long UMIs. If a long UMI is chosen, UMI diversity is high and the number of UMIs is much greater than the number of molecules of the major allele, resulting in a small UMI collision rate. In addition, if UMI diversity is high, sequencing errors that affect the UMI can be corrected. For example, if all UMIs present in a sequenced population are at least two nucleotide changes from each other, sequenced UMIs that differ by only one nucleotide change can be merged (Smith *et al*., 2017).

However, we note that excessively long UMI lengths chosen without consideration for these concerns pose several complications: First, a greater number of sequencer cycles spent on reading UMIs necessitates a shorter read of the actual target sequence. Second, there is some speculation that long UMIs could interfere with primer sequence binding— by preferentially hybridizing to DNA with complementary sequence, or sterically hindering proper adapter function at the target site. Third, given that each basepair has an equal probability of sequencing error, longer UMI sequences are more likely to accumulate sequencing errors, which could result in ambiguity about whether sequences with similar UMIs result from sequencing errors or are unique molecular events.

To quantify the effects of UMI length on UMI collisions, we have derived an equation, Equation (4), that allows us to compute the probability of observing no UMI collisions as a function of UMI length, number of reads, and the number and proportions of alleles. The calculation of this probability for the worst case is implemented in our software and can be calculated using the `AmpUMI Collision` command. In some cases, a small number of UMI collisions may be tolerable as long as the final allelic proportions are within a certain error. In this case, Equation (13) can be used to determine the total allelic fraction distortion for a given UMI length and the true underlying allelic composition. The calculation of allelic distortion can be computed using the `AmpUMI Distortion` command. AmpUMI can also be used to determine the minimum UMI length required to have a probability of observing a UMI collision below a given threshold, or to produce an allelic distortion below a given percentage. These commands can be accessed using the `AmpUMI Collision or AmpUMI Distortion` command as before, except that the desired cutoff is given as a parameter instead of the UMI length.

We note that there is a difference between the number of reads and the number of DNA fragments being sampled. While it is relatively easy to predict the number of reads that will be produced by a sequencing run based on the machine type and the abundance of the sample among those being sequenced, it is impossible to predict the allele frequency of the most frequent allele. With this in mind, we suggest that for experiments for which the major allele frequency is not known, a suitable UMI length be chosen based on the case that the amplicon sequencing yields a single allele. In this scenario, if we assume that we have *n* reads, we can use the `AmpUMI Collision` command or Equation (6) to determine the minimum UMI length *k* for observing no collisions with a given probability *p*.

### 2.2 AmpUMI: Removing PCR duplicates from amplicon sequencing

Once the proper UMI length has been designed and the experiment has been sequenced, the raw FASTQ output from the sequencer needs to be prepared for downstream analysis. In addition to aiding in UMI design, AmpUMI is a flexible and efficient software that can be used to perform preparation of raw FASTQ files. FASTQ files contain the sequence of the UMI and DNA fragment, as well as the sequencing quality for each read. Unprocessed FASTQ files cannot be used in downstream processing steps because the data still contains PCR duplicates, and the UMI sequence must be separated from the sequence of the DNA fragment. AmpUMI performs four major preprocessing functions: assessing the quality of UMIs, flexible trimming of the UMI and adapter sequences, performing sequencing error correction, and finally removing reads resulting from PCR duplicates.

First, AmpUMI assesses the quality of UMIs. In this step, the diversity of UMIs is measured. If one or a few UMIs are significantly overrepresented in the library, this could signal an error in UMI synthesis or bias in library amplification. In addition, we propose a measure for estimating whether the UMI complexity was sufficient for the diversity of amplicon sequences. We report the frequency of reads with the same UMI that have different amplicon sequences. A high frequency of these types of reads would result from an experiment with too little UMI complexity, or from a highly error-prone sequencing run.

Next, the UMIs and adapter sequences are trimmed from the FASTQ sequences. AmpUMI allows users to specify a regular expression to flexibly specify the UMI and adapter sequences. In the specification of this regular expression, the locations of bases that should be counted as UMIs are set by the user as the letter *I*, while the sequencing adapter is specified using the regular *A*, *T*, *C*, and *G* letters. Using this regular expression, the UMI and adapter are identified and trimmed from each read. The UMI for each sequence is added to the sequence name. Reads for which the regular expression cannot be found (e.g., due to sequencing errors in the sequencing adapter or length modifications of the UMI) are discarded.

After the reads are trimmed, sequencing error correction and PCR deduplication are performed. Two functions for error correction are used in the deduplication process. The first type of error correction is designed for experiments in which the number of UMIs is much greater than the number of input molecules and multiple rounds of PCR have been performed. This experimental design will yield multiple copies of each UMI-molecule pair, and sequencing errors will be apparent as rare mutations in the amplicon sequence. During deduplication, reads will be grouped by UMI, and the most frequent amplicon sequence will be accepted as the consensus sequence and the other sequences will be assumed to come from sequencing error and will be discarded. The second type of error correction is optional and can be used to correct errors in the UMI sequence by filtering rare UMI-amplicon sequence pairs that could arise from sequencing errors. First, FASTQ sequences with the same UMI and amplicon sequence are grouped. Only pairs of UMI-amplicon sequence that have been seen multiple times (a parameter specified by the user) are used for downstream analysis, while the remainder of rare UMI-amplicon sequence pairs are discarded. As this parameter is increased, the confidence in the UMI and amplicon sequence is increased, but the quantification accuracy of allele distribution is decreased, especially for rare alleles.

## 3 Methods

### 3.1 Problem formulation and assumptions

We are interested in determining the probability of no collisions of any two different alleles associated with the same UMI. In what follows, we make a few motivated assumptions to derive a closed-form expression for our probability of interest.

Let *X* be finite set of *M* (= |*X*|) DNA fragments partitioned into *I* types (i.e. alleles) and let *Y* be a finite set of *U* (= |*Y*|) UMI primers partitioned into *J* UMIs (i.e. barcodes), such that fragments or UMI primers within each class are indistinguishable from each other. Here, the number of UMIs *J* equals the number of the four nucleotides (A, G, C, T) raised to the power of the UMI’s sequence length *K*. That is, *J* = 4^*K*^, where the value of *K* usually ranges from 8 to 12bp. Accordingly, the two sets of interest can be partitioned into *X* = {*X*_1_, …, *X*_*i*_, …, *X*_*I*_} and *Y* = {*Y*_1_, ⋅, *Y*_*j*_, ⋅, *Y*_*J*_} where |*X*_*i*_| = *M*_*i*_ for *i* = 1, …, *I* and |*Y*_*j*_| = *U*_*j*_ for *j* = 1, …, *J*.

#### Assumption 1 (Sampling with Replacement)

Given that the total number of UMI primers *U* is much greater than the number of unique UMIs *J* and the total number of DNA fragments *M* is also much greater than the number of alleles *I*, we can assume that both the UMI primers and DNA fragments are sampled with replacement. Note that the number of reads (samples) *n*, which we introduce later, is also assumed to be much less than *U*. For the UMIs, the assumption is motivated by the observation that in the PCR amplification step, there are typically 3 × 10^11^ UMI-labeled primers in 0.5 pmol of solution and so for UMIs of length *K* = 10, we would, on average, have approximately 286,100 copies for each UMI.

Assumption (1) implies that the probability of sampling either allele *i* or UMI *j* is given by the categorical distribution (a generalization of the Bernoulli distribution to multiple categories). That is

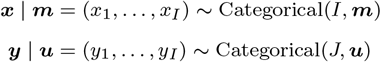

where ***x*** ∈ {1, 2,…, *I*}, ***m*** = (*m*_1_,…, *m*_*I*_), ***y*** ∈ {1, 2,…, *J*}, and ***u*** = (*u*_1_,…, *u*_*J*_). The proportion 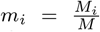 represents the probability of observing allele *i* such that 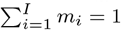. Similarly, the proportion 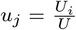 represents the probability of observing UMI *j* such that 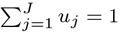.

Now, the total number of reads (samples) *n* equals the number of UMI-allele pairs—of which there are *Q* = |Ω| = *I* ×*J* unique combinations. To determine the distribution over unique UMI-allele pairs, we first define a joint sample space Ω = {(1, 1), (1, 2),…, (*i, j*),… (*I, J*)} to enumerate all *Q* possible pairing outcomes and we let the random variable ***z*** be defined over Ω.

#### Assumption 2 (Independence)

Since there is no indication that a given UMI has a preference for any allele, we can assume that the pairing process is unbiased such that sampling ***x*** is independent of sampling ***y***.

Assumption (2) implies that the probability mass function for ***z*** is given by a multinomial distribution with support ***z*** = {(*z*_1_,…, *z*_*Q*_) ∈ ℕ^*Q*^ |*z*_1_ + … + *z*_*Q*_ = *n*} where *n* is the number of samples and ***p*** = ***m*** ⨂ ***u*** = [*m*_1_*u*_1_,…, *m*_1_*u*_*j*_,… *m*_*i*_*u*_*j*_,… *m*_*I*_*u*_*J*_] is the probability vector for the *Q* possible pairings, where ⨂ is the Kronecker product. That is,

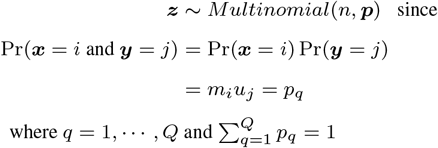

##### Example

For the case of two alleles *I* = 2, three UMIs *J* = 3, and three observations (i.e. reads) *n* = 3, the probability of observing two reads from Allele 1 paired with UMI 1 and one read from Allele 2 paired with UMI 3 is

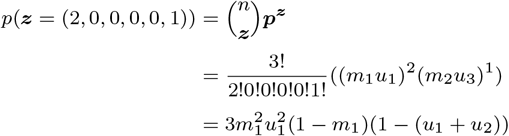

To derive the probability of obtaining no collisions, we note that the multinomial distribution for a sample size *n* can be viewed geometrically as the normalized *n*’th component of a *Q−*dimensional Pascal’s simplex, ∧^*Q*^, whose *n*’th component, 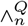, consists of the coefficients of the multinomial expansion of the polynomial 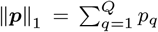 raised to the power *n*. In particular, according to the multinomial theorem, we have

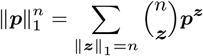

where ***p** ∈* [0, 1]^*Q*^, 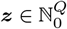, *n* ∈ ℕ_0_, *Q* ∈ ℕ, and ||***p***||_1_ = 1.

Therefore, the number of multinomial coefficients is equal to the number of coefficients in the (*Q −* 1) simplex 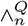 and is given by the (*n* + 1)th simplicial (*Q −* 1)-polytopic number

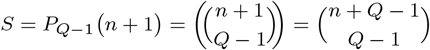

Each of the *S* components of the multinomial expansion represents the probability of a particular counts outcome event (i.e.***z*** = *z*), which is given by the multinomial distribution

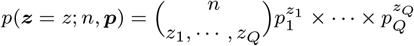

We are interested in a subset of the *S* events that correspond to having no collisions. We denote the set of no collisions by

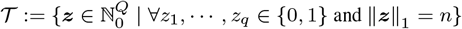

Note that the cardinality of 𝒯 is equal to the number of ways a total of *n* 1s are selected from a group of *n* 1s and (*Q − n*) 0s. More specifically,

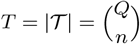

Now since for all T events, the product of the terms in denominator of multinomial coefficient is equal to 1, we can simplify the expression for the probability of observing ***z***\* ∈ 𝒯 as such

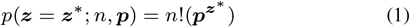

Note that when *n > Q*, the probability of observing any of the *T* no-collision events is zero, otherwise when *n ≤ Q* the probability of observing no collision can be expressed concisely as

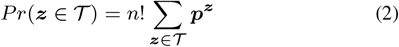

#### Assumption 3 (Equal Proportions of UMIs)

Since the relative abundance of UMIs can be controlled during UMI synthesis, we can assume that the proportion of UMIs to be uniform such that 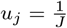 for all *j, j* = 1,… *J*.

Assumption (3) allows us to further simplify Equation (1) as such

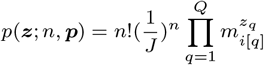

and so the probability of observing any of the *T* no-collision events, Equation (2), becomes

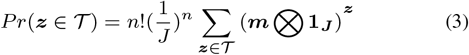

Computing the probability given by Equation (3) is intractable for large values of *T*. Nonetheless, the expression can further be simplified if we assume the following.

#### Assumption 4 (Upper Limit on the Number of Reads)

When the UMI sequence length *K* is 10, *J* already exceeds 1 million. For some experiments then, we can assume that the number of reads is less than or equal to the number of UMIs *n* ≤ *J*

Using Assumption (4), we can partition the set 𝒯 into *E* equivalent classes, essentially reducing the set into an *I*-dimensional *discrete simplex*

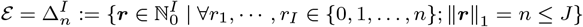

with cardinality

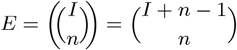

such that

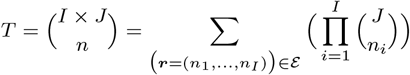

so now the probability of observing any of the *T* no-collision events is given by this general formula.

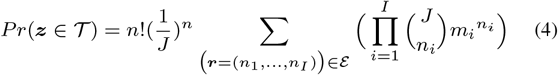

### 3.2 Special cases

#### Case 1 (One Allele (*I* = 1, Worst Case))

When *I* = 1, the discrete simplex *ε* has just a single component. Subsequently, Equation (4) reduces to

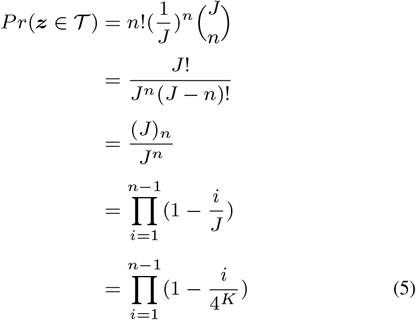

To find the minimum UMI length *k\** for a given sample size *n\** such the probability of no collisions is above a certain threshold *p\**, we first need to compute the quantity given by Equation (5) for a given *n\** for each UMI length up to an upper bound *k*_*max*_ (say 20). Let us denote the vector of computed probabilities by

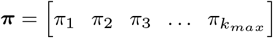

Note the elements of ***π*** corresponding to values of *K* for which *n <* 4^*K*^ are zero and do not require computing. The minimum UMI length *k\** can then be formally written as

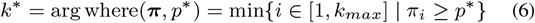

where arg where(***π**, p\**) gives back the index of the first vector’s element whose value exceeds *p\**. Note that since the value of *π*_*i*_ increases as *i → k*_*max*_, we only need to return the index of the first element exceeding the threshold.

#### Case 2 (Equal Proportion of Alleles)

Equations (3) and (4) have the simplest form when ***m*** is a vector of equal probabilities. That is, when 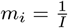 for all *i*, *i* = 1, … *I*. The probability of observing any of the *T* no-collision events simplifies as such.

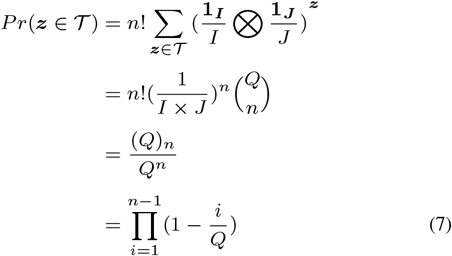

where (*Q*)_*n*_ denotes the falling factorial. Note that the probability of no collision approaches 0 very rapidly as *n* goes from 1 to *Q*. Also, note that when *n > Q*, the probability of collision is 1 for any *Q*.

Note that whereas the number of multinomial coefficients is given by *S* for the set of all events and *T* for the set of no-collision events, the sum of the coefficients is given by *A* = *Q*^*n*^ for the set of all events and by

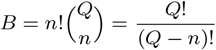

for the set of no collisions. Alternatively, the probability can be derived as a ratio of B over A

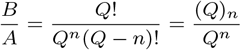

#### Case 3 (Two Alleles (*I* = 2))

When *I* = 2, the discrete simplex *ε* has has *n* components. Subsequently, Equation (4) reduces to

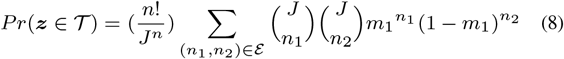

where the values of ***r*** = (*n*_1_, *n*_2_) vary over the anti-diagonal indices of an *n × n* matrix and the discrete simplex is given by

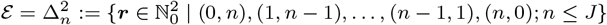

### 3.3 Effects of Collisions on Allelic Diversity

Let ***y*** = (*y*_11_,…, *y*_1*J*_,…, *y*_*I*1_,…, *y*_*IJ*_) denote the *q*-dimensional vector of deduplicated reads for all UMI-allele pairing such that *y*_*ij*_ ∈ {0, 1} refers to the read corresponding to the *i*th allele and *j*th UMI. The model, where an indicator function 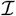 thresholds each element of the unobservable random vector ***z*** into an observable binary variable *y*_*ij*_.

That is,

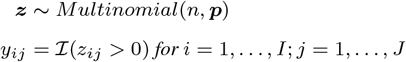

where 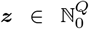, ||***z***||_1_ = *n*, ***p*** ∈ [0, 1]^*Q*^, ||***p***||_1_ = 1, *n* ∈ ℕ_0_,, and *Q* = (*I × J*) *∈* ℕ

#### Expected Number of Deduplicated Reads

We are interested in the expected number of deduplicated reads (i.e. one-valued elements) in each of the *I* allele partitions of ***y***. We begin by letting **M** denote the *I ×*(*I ×J*) transformation matrix that gives the *I*-dimensional vector of deduplicated read counts for each allele. That is,

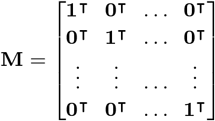

Accordingly, the vector of expected deduplicated read counts ***c*** can be obtained as such

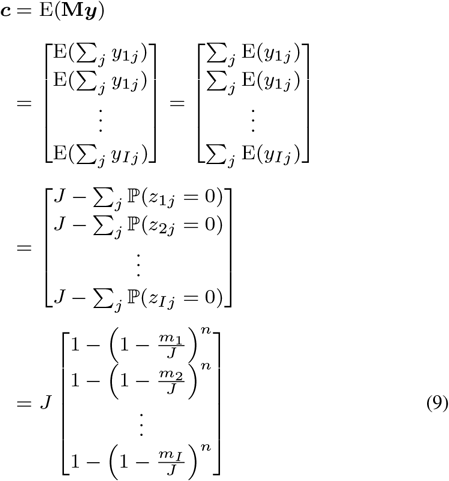

The penultimate step was obtained by observing that the expectation of an event defined by an indicator function is equal to the probability of that event. That is

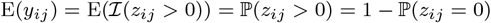

The final step was obtained by noting that since the marginal distribution of each element of the multinomial ***z*** is binomial, namely,

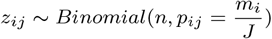

then the probabilities of interest are just given as such.

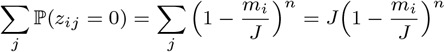

##### Expected Number of Collisions

The expected number of collisions ***n***_*c*_ can be computed by noting that the number of collisions for each UMI-allele pairing is given by the vector ***x*** = ***z** − **y*** and so the vector of expected number of collisions for all the alleles is given by

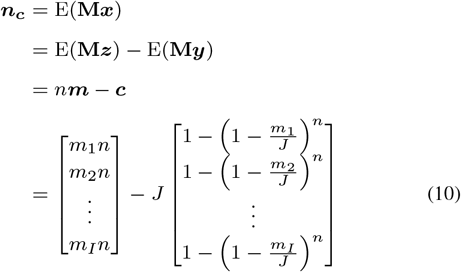

Furthermore, the expected total number of collisions collapsed over the alleles is given by

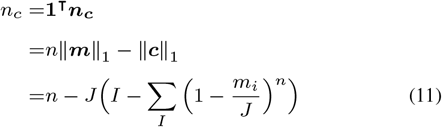

##### Allelic Fraction Distortion

The expected allelic fraction distortion is simply the difference of the two normalized components of ***n***_***c***_. More specifically, since the elements ***c*** are all non-negative, we can formally define and simplify the quantity as such.

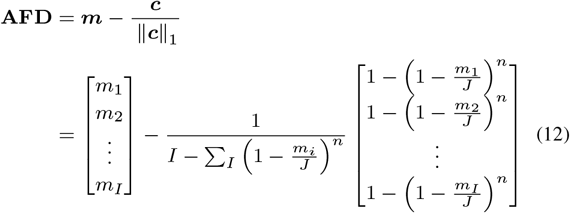

which can be reduced to a scalar quantity, the expected total allelic fraction distortion, which we abbreviate as **TAFD** and can be obtained as such.

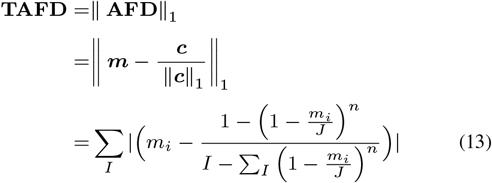

## 4 Results

### 4.1 Allelic fraction distortion resulting from inadequate UMI length

One of the purposes of performing amplicon sequencing is to accurately estimate the frequency of each allele variant in the population. However, we show here that the observed allelic fraction of each variant can be distorted if the UMI length is too short and the UMI complexity is too low. In the case in which allele variants are present at roughly the same proportion in the population (*m*_1_ *≈ m*_2_ *≈ m*_3_ *≈ m*_*I*_) this distortion will be small, as UMI-molecule collisions will affect each molecule type equally. However, as the proportion of allele variants becomes more imbalanced (e.g., *m*_1_ ≫ *m*_2_ ≫ *m*_3_ …), after deduplication using UMIs the observed frequency of rare alleles will increase, and the observed frequency of frequent alleles will be lower than the actual frequency in the population. Intuitively, this is because collisions are more frequent in the more frequent alleles, where a larger UMI diversity is required to avoid UMI-molecule collisions. Rare allele with few reads are less likely to exhaust the available unique UMIs.

As an example, we simulated biological library preparations with *n* =100,000 reads from a population with an underlying allelic diversity defined as ***m*** = (0.1, 0.1, 0.3, 0.5). In each simulation, alleles were paired with a UMI. Samples were simulated using UMIs of length between 1 and 18bp long. After the simulated sample was assembled, reads with the same UMI and allele were deduplicated, leaving only one unique UMI-allele combination in the sample. The proportion of each allele was reported. As seen in Figure 2, if the UMI is too short and UMI complexity is too low then the proportion of each allele, including rare alleles will be approximately equal. This has the effect of overestimating the frequency of rare alleles in the population, while underestimating the frequency of abundant alleles as shown on the left side of Figure 2. With sufficient UMI diversity (right side of Figure 2), the true, underlying allele frequencies are recapitulated in the simulated, deduplicated populations. As expected, the allelic frequencies produced by our model are in perfect agreement with those obtained with computationally expensive simulations.

**Fig. 2.**
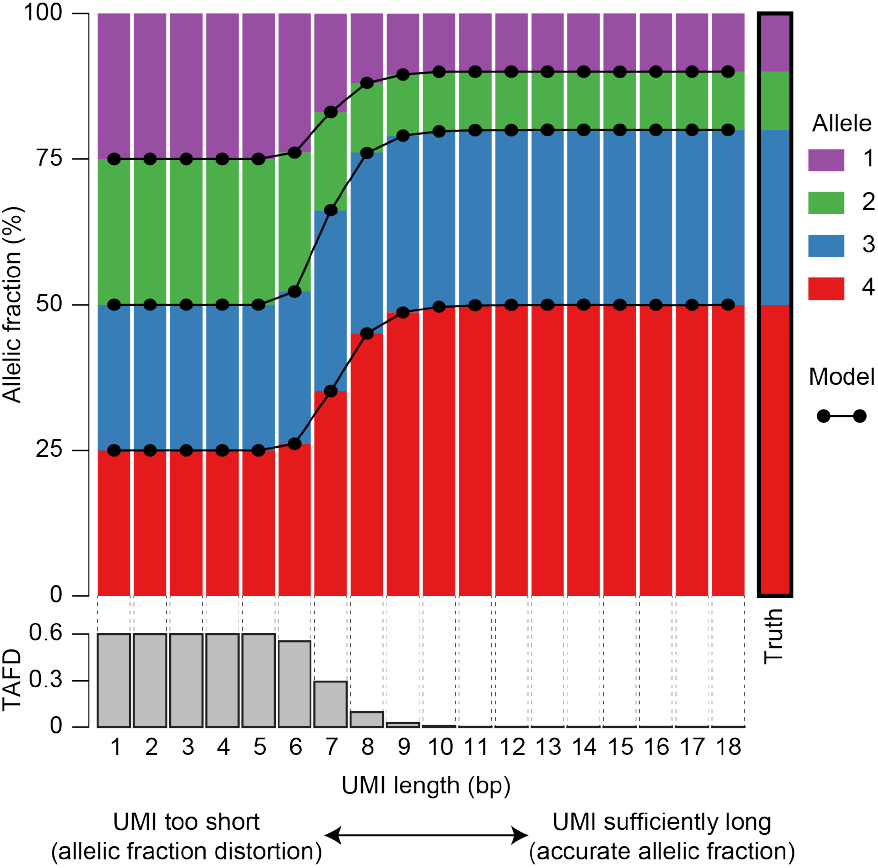
Association between distortion of allelic frequency and UMI length. Colored bars show simulated allelic fractions of four alleles after deduplication of reads with the same UMI and allele. Simulated samples consisted of *n* =100,000 reads and were drawn from a population with an allelic diversity given by ***m*** = (0.1, 0.1, 0.3, 0.5). Reads were generated using using UMI of length between 1 and 18bp long. For each UMI length, the average simulation proportion of each allele is shown after removing UMI-allele collisions. 100 samples were simulated for each UMI length. The right column marked ‘Truth’ shows the underlying allelic diversity from which the simulated samples were drawn. Dots connected by lines show the predicted allele frequency given our model (Equation (12)) and are in complete concordance with the simulation results. The gray histogram at the bottom of the plot shows the total allelic fraction distortion (TAFD) (Equation (13)) for each UMI length.

We then considered the number of collisions in our simulated samples (Figure 3). The number of collisions is defined as the number of simulated reads (UMI-molecule combination) that had been previously observed in the simulation. For example, We simulated 100,000 reads with a UMI of length 1 (4 possible UMIs), and measured the collision rate as 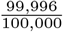. We note that the accuracy of recapitulating the actual allele fraction as shown in Figure 2 is mirrored in the collision rate shown in Figure 3.

**Fig. 3.**
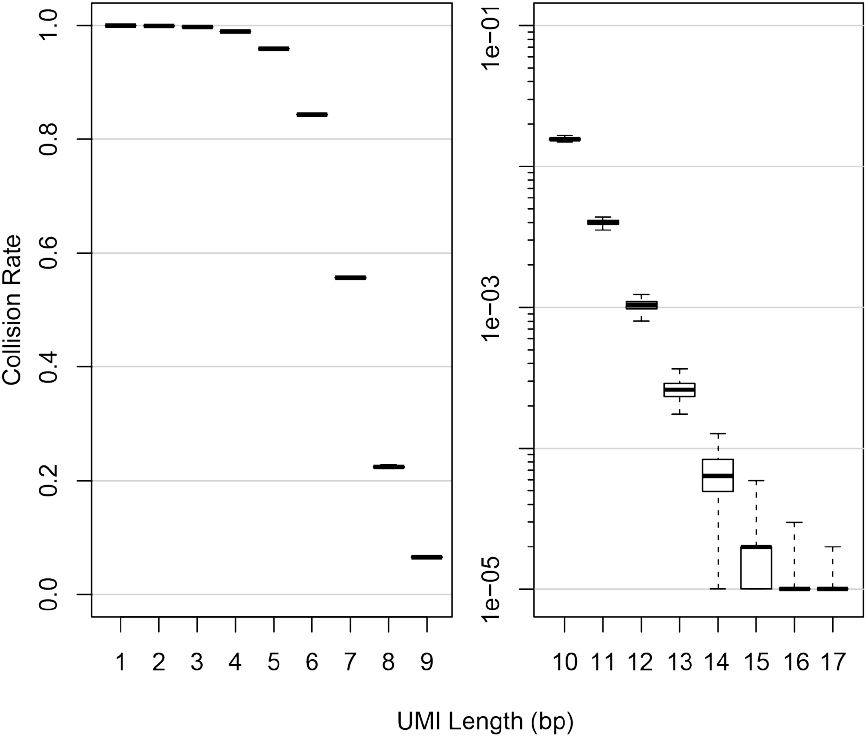
Distribution of collisions in simulated populations. The number of collisions in each simulated sample used in Figure 2 were aggregated by UMI length. Boxplots show the median (thick line), interquartile range (box), and the range of the data (whiskers). The count of collisions is defined as the number of simulated reads (UMI-molecule combination) that had already been observed in the simulation.

### 4.2 Probability of observing no UMI collisions

Next, we address the problem of finding the probability of observing no UMI collisions. Equation (5) describes the probability of observing no collisions in the worst case scenario in which we are most likely to observe a UMI collision—when only one allele is present in our DNA fragment pool. Figure 4 shows the probability of having no UMI collisions for a variety of UMI lengths. When we construct 100,000 reads consisting of a random UMI and a DNA fragment, an 18bp UMI yields a sample with no collisions with probability p=0.93, while shorter UMIs are less likely to yield a sample with no collisions.

**Fig. 4.**
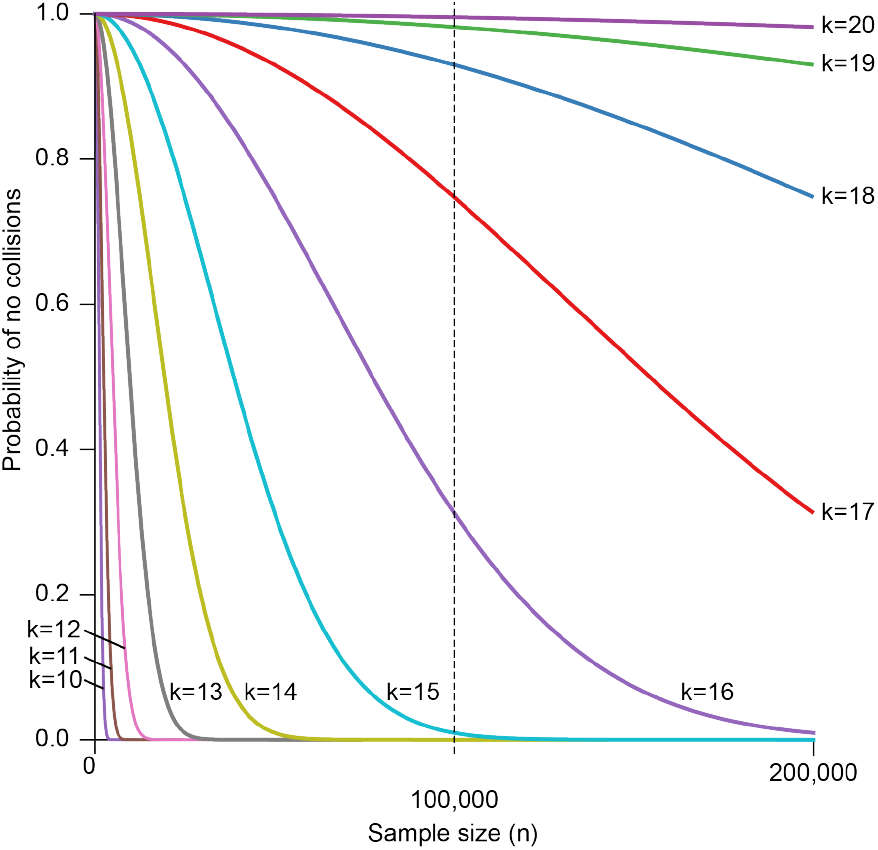
Probability of having no UMI collisions for Case 2 (worst case scenario): The probability of no collision as a function of sample size *n* for 5 consecutive values of UMI lengths, *k* = 14,…, 18 such that *J* = 4^*k*^ ≤ *n* (colored curves). The vertical dotted line shows the n=100,000 sample size referenced in Figures 2 and 5.

#### 4.3 Rates of no UMI collisions in simulated samples

We wanted to validate our model with simulated data, so we simulated samples of size *n* =100000 with a UMI of length between 6bp and 17bp long. For each simulation sample, we simulated a set of DNA fragments consisting of five different alleles. The fractional presence of each allele was randomly selected for each simulation sample to explore collision rates in cases without prior knowledge on allelic composition of a sample. DNA fragments were paired with UMIs and the number of UMI collisions was recorded. A total of 1000 simulations were performed for each UMI length. Figure 5 shows the average percent of simulated samples in which no UMI collisions were recorded. The red line in this figure shows the predicted probability for the worst case in which there is only one type of molecule. As expected, the model-predicted worst case (red line) is always below the average of all simulated samples (blue lines) for every UMI length.

**Fig. 5.**
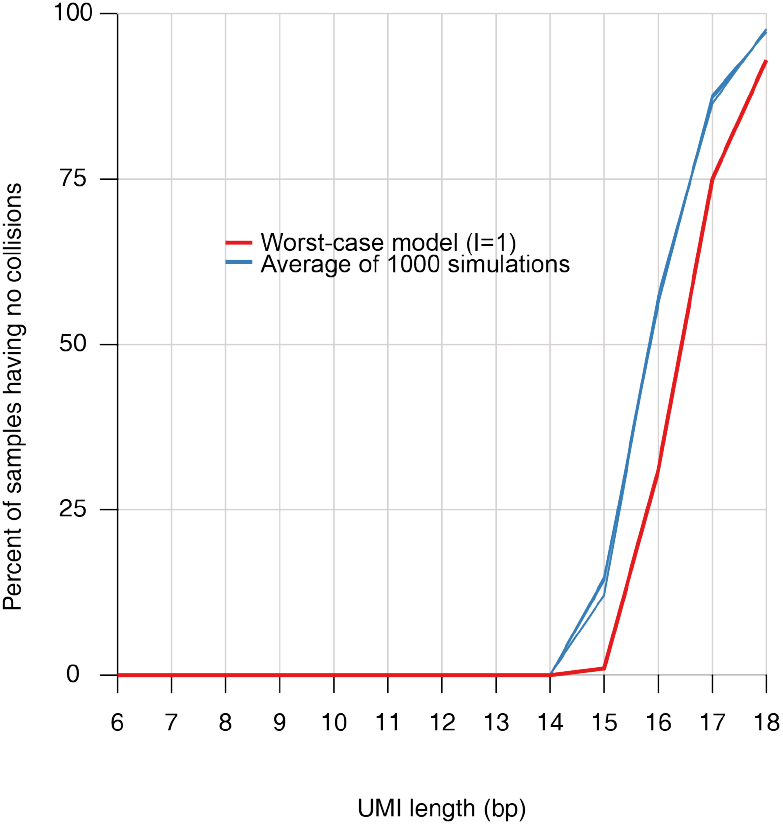
Probability of having no UMI collisions in simulated samples. We simulated samples of size *n* =100000, with DNA fragments randomly selected from a set containing 5 unique fragments each with a random fraction of presence in the sample. Simulated DNA fragments were paired with a given set of UMIs, and the rate of UMI collisions were measured. The average percent of all 1000 simulated samples having no collisions is shown with the blue line. 3 simulations were carried out with 1000 samples each. The red reference line is computed by our model, and shows the values in Figure 4.

We were surprised to observe that the length of UMI required to achieve no UMI collisions was so long—with a minimum of 18bp long for a sample of 100,000 DNA fragments. This indicates that for experiments performed with this number of fragments and a shorter UMI (length 8bp or length 12bp), a small number of undetectable PCR duplicates are likely present in these libraries.

We note that the probability of having no collisions may be somewhat conservative for some experimentalists who may have a certain tolerance for an allowable number of collisions in an amplicon sequencing experiment. In this case, Equation (13) or the `AmpUMI Distortion` command can be used to determine the required UMI length after selecting a tolerable allelic distortion for their sequencing application.

#### 4.4 AmpUMI validation on experimental data

To validate our deduplication software, we analyzed a publicly-available dataset of deep amplicon sequencing (Kou *et al*., 2016). This dataset was created to measure PCR amplification and sequencing errors, and utilized a 22bp UMI. The amplicon sequence was verified by Sanger sequencing to match the reference sequence. We tested to see whether AmpUMI could correctly remove erroneous, non-reference reads. Table 1 shows data for the unprocessed data and after the first and second steps of data processing using AmpUMI. In Step 1, reads with the same UMI are deduplicated, and only the most-frequent allele is kept for each UMI. In Step 2, UMI-sequence pairs that appear less than 10 times are filtered out. Table 1 shows the number of total reads, the number of unique alleles, and the percentage of reads that represent the true, unmodified, reference allele. Although only 95.77% of reads in the unprocessed sample match the unmodified allele sequence (the remainder of reads contain PCR and sequencing artifacts), after both steps of processing by AmpUMI, 99.19% of reads match the unmodified allele, showing a substantial reduction in contaminating reads resulting from PCR or sequencing artifacts.

**Table 1.** Number of reads, alleles, and the percent of reads matching the reference allele that are present before and after AmpUMI analysis of a dataset of deep amplicon sequencing (Kou et al., 2016).

|  | No. Reads | No. Alleles | Pct. Ref Allele |
| --- | --- | --- | --- |
| Unprocessed | 6,440,216 | 20,760 | 95.77 |
| Deduplicated - Step 1 | 426,447 | 2,524 | 97.05 |
| Deduplicated - Step 2 | 269,564 | 176 | 99.19 |

## 5 Discussion

In this study, we address the problem of UMI design and outline a statistical framework for calculating the probability of collisions of different molecules having the same UMI. We also introduce AmpUMI, a flexible software for analyzing next-generation sequencing from experiments using UMIs and for preparing data for downstream analysis.

Our statistical framework allows researchers to calculate the probability of having no collisions for a given experiment, and can be used to determine an acceptable number of UMIs in order to reduce the probability of observing a collision to an acceptable threshold as determined by the user. In cases where a small number of UMI collisions are acceptable, our framework can also be used to estimate the allelic distortion resulting from these collisions, and an appropriate UMI length can be chosen to keep the allelic distortion below a specified threshold.

We show that AmpUMI is a valuable tool for analyzing amplicon sequencing using UMIs. AmpUMI first allows users to assess the quality of UMIs and to determine whether the UMI pool had adequate diversity to prevent UMI collisions. Next, flexible trimming of the UMI and adapter sequences allows researchers to adapt the software to their specific UMI and sequencing design. AmpUMI can also performing sequencing error correction to reduce noise and mitigate sequencing errors. Finally PCR duplicate reads are removed from the sample FASTQ for easy integration into downstream analysis.

Although many analysis tools exist for deduplicating next-generation sequencing data, none are designed to use UMI information for deep amplicon sequencing in cases where preservation of allelic frequency is important. AmpUMI will fill a gap in the available analysis tools that are currently available to the community. We anticipate that this tool will be useful in a variety of applications. For example, in the genome editing community, much effort is being placed on the quantification and characterization of CRISPR activity in the genome and the identification of off-targets (Tsai *et al*., 2015; Kim *et al*., 2015; Tsai *et al*., 2017). The gold standard for analyzing mutations at targets is by deep amplicon sequencing of the target which allows for quantification of editing frequency and analyzing genetic editing events (Pinello *et al*., 2016). After performing genomic editing, the rate of modified reads is compared to the rate of unmodified reads to determine editing efficiency at that location. Because a large percent of the resulting amplicon sequences are the unmodified sequence, simply removing duplicate sequences would greatly distort the measure of editing frequency, and it is impossible to tell which sequence are arising from PCR duplication or unique molecules. Sequencing strategies that use UMIs could greatly improve the accuracy and confidence in this measure.

When performing amplicon sequencing on a sample with unknown allele frequency, we recommend using the collision rate calculated for a sample with only one allele, and acknowledge that this will yield a very conservative estimate of minimum UMI length. In a laboratory setting, this conservative UMI length could be used in a pilot study sequenced at low sequencing depth to better understand the underlying allele distribution, after which a more practical UMI length can be established for full-scale sequencing analysis.

In conclusion, we provide AmpUMI as an open source tool that we hope will benefit the scientific community, particularly in the design of appropriate UMIs and in the processing of sequencing reads produced by amplicon sequencing using UMIs. AmpUMI is open-source and freely available at http://github.com/pinellolab/AmpUMI.

## Author Contributions

R.F. and K.C. conceived the methodology; K.C. developed the software; R.F. developed the statistical formalism and worked out the mathematical derivations; R.F. and K.C created the visualization plots; K.C., R.F., D.E.B., and L.P. wrote, reviewed, and contributed to the final version of the manuscript; K.C., R.F., D.E.B., and L.P. conceived the ideas in the paper. L.P. supervised the project.

